# Associations between amygdala connectivity and experienced discrimination in children

**DOI:** 10.64898/2025.12.22.695992

**Authors:** Nicholas J. Goode, Eric Potter, Meerab Rasheed, Mariela Camacho, Rinko Higashi, Linh Nguyen, Shubangini Shah, Oscar Miranda-Dominguez, Robert Hermosillo, Anthony Juliano, Aya Cheaito, Elina Thomas

## Abstract

Discrimination is a chronic stressor linked to adverse health outcomes, particularly in racial and ethnic minorities. Understanding associations between early discrimination and the brain in childhood may help identify mechanisms through which discrimination impacts future health. Data from 4512 children (ages 9-11) and a subsample of Black, Indigenous, and People of Color (BIPOC; N = 1567) from the Adolescent Brain and Cognitive Development (ABCD) Study® was used to create linear mixed-effects models that evaluated associations between Perceived Discrimination (PD) and amygdala resting-state fMRI connectivity (AC) to the salience network (SN), default mode network (DMN), and thalamus. PD was measured using the youth self-reported PD Scale. Results indicated that greater PD significantly predicted greater AC to the right thalamus in our full sample. In secondary analyses, environmental and behavioral factors were evaluated as potential moderators for associations significant at least at a trend level in both our full sample and BIPOC subsample. In our BIPOC subsample, traumatic events experienced moderated the relationship between PD and AC to the anterior cingulate cortex (ACC; SN), such that greater traumatic experiences predicted stronger positive associations between PD and this connection. Results suggest PD impacts neural connections in early life, highlighting the need to consider the impact of discrimination on risk for psychopathology.

## 1. Introduction

Discrimination, unfair or prejudicial treatment based on characteristics such as race, ethnicity, gender, or sexuality (1, 2), has been recognized as a public health crisis in the United States (3, 4). As a chronic stressor, discrimination has been shown to increase allostatic load (5), which results in poor health outcomes, including increased inflammation (6), compromised immunity (7), poor cardiovascular health (2, 8, 9), and alterations in sleep (10, 11) and stress regulation (12–15). Discriminatory experiences are also associated with an increased risk for psychopathology. In particular, increased experiences are linked to increased internalizing and externalizing behaviors (9, 16), suicidality (9, 17), and post-traumatic stress symptoms (18–20). In the United States, Black, Indigenous, and People of Color (BIPOC) face disproportionate levels of discrimination (21, 22), which may lead to greater impacts on mental (23–25) and physical (26) health.

Awareness of experienced discrimination typically begins before the age of 10, and increases with age, allowing for the examination of the impact of these experiences in childhood (27–29). Prior literature indicates that experiences of discrimination at this time point increase risk for poor health through disruptions in the development of the brain and other biological systems (30, 31). As in adulthood, sleep (32, 33) and stress dysregulation (32) also occur in response to these experiences, further impacting health. Psychological health is particularly affected, as increased externalizing and internalizing behaviors (34–38), feelings of hopelessness (38), and suicidality (39, 40) are observed with increased exposure to discrimination. Experienced discrimination may also increase the impact of trauma on psychological outcomes (41) at this age, though the biological mechanisms that support this are poorly understood. Despite research on the adverse effects of discrimination (30–32), little is known about how early experiences of discrimination affect neural connectivity in childhood. Understanding these associations may help clinicians and researchers choose and identify interventions (42, 43) that target specific brain connections to reduce the impact of these experiences.

In adulthood, discriminatory experiences have been linked to harmful structural changes in the brain, including reduced overall volume (7; 44) and white matter tract integrity (45, 46). Several regions and networks involved in fear learning (47, 48), emotion regulation (49), and memory and perception (50, 51) appear to be consistently impacted. These regions include the amygdala (52–55), insula (55–57), anterior cingulate cortex (58), and hippocampus (59). These particular regions act as nodes (60) in the salience (SN) and default mode networks (DMN), which also show changes in connectivity patterns in response to experiences of discrimination (56, 57, 61). Across the literature, amygdala connectivity is consistently impacted (55, 62, 63), particularly to nodes in the SN (52, 64) and DMN (56, 65) and thalamus (52, 55, 63). While associations between these connections and discrimination have been established in adults, these relationships have not been evaluated in childhood.

In BIPOC adults, heightened levels of discriminatory experiences (21, 22) may cause amygdala connections associated with discrimination to be further impacted. This may explain the intergenerational effects on amygdala connectivity (62, 63) and additional impacts on mental (23–25) and physical health (26) that have been observed in this population. Thus, it may be beneficial to consider BIPOC populations separately when examining associations between these amygdala connections and experienced discrimination in childhood.

While much is known about how experienced discrimination affects the brain in adulthood, less is known about its impacts on the brain in childhood. Existing literature shows that these experiences impact the brain broadly, slowing whole-brain growth (66) and affecting similar structures such as the hippocampus and amygdala (67). In particular, greater experiences accelerate the maturation of these structures (67) and affect hippocampal volume and reactivity (68, 69;), and amygdala connectivity (67) and volume (68, 70). While these associations appear to be in line with findings in adulthood, amygdala connectivity to regions most consistently impacted in adulthood (thalamus, DMN nodes, and SN nodes) has not been evaluated. Given the amygdala’s involvement in fear processing (48) and stress regulation across development (71–73), further examination of how discrimination impacts these amygdala connections in childhood is warranted.

Associations between experienced discrimination and the brain in childhood may be influenced by behavioral and environmental factors. Internalizing and externalizing behaviors likely exert an impact. The relationship between discrimination and these behaviors appears to be cyclical. The presence of internalizing and externalizing behaviors may increase the influence of discrimination on the brain, while experiences of discrimination may increase engagement in these behaviors (74, 75). Both internalizing and externalizing behaviors, like discrimination, are associated with changes in amygdala volume and connectivity to SN and DMN nodes implicated in emotion regulation (e.g., the PFC and ACC (76, 77)), which may lead to increased reactivity to threat (78, 79), potentially amplifying the effect of discrimination on the brain.

Some environmental factors may mitigate the impact of discrimination, such as the child’s school environment and engagement in prosocial behavior. A school environment that promotes feelings of safety, teacher support, and opportunities for involvement may facilitate resilience to the effects of these experiences (80–82). However, in instances where exposure to discrimination is increased in this setting, resilience may be reduced. Prosocial behaviors may also protect against the negative impacts of discrimination (83–85). These behaviors and the school environment can facilitate social connection (86), which improves brain (87, 88) and mental health outcomes (85). Social connection between peers belonging to similar racial, ethnic, or gender groups may be particularly impactful, as increasing one’s sense of belonging and identity has been shown to improve coping strategies and outcomes (89–93). A supportive school environment can foster these connections (82). Given these prior associations, the supportive nature of the school environment and engagement in prosocial behaviors should be considered when evaluating the impact of discrimination on the brain.

Other environmental factors, like family conflict, English language proficiency, and experienced traumatic events, may also influence associations between discrimination and the brain. Increased family conflict is linked to poor mental health (94, 95), which amplifies the negative impact of discrimination on health overall (96, 97) and likely impacts the brain. Similarly, a lack of proficiency in the English language increases the amount of discrimination one experiences (98–100), resulting in poor mental health outcomes, such as increased experienced distress, depression, and anxiety (101, 102). Additionally, experienced traumatic events may increase sensitivity to the effects of discrimination by overactivation of stress-related brain regions like the amygdala, hippocampus, and PFC (65, 89; 103), potentially exacerbating its impact on the brain. As a result, those who have experienced both trauma and discrimination often show greater PTSD symptoms like hyperarousal and intrusive symptoms of stress (89, 103–105). Disparities in experienced adversity between BIPOC and non-BIPOC children may further explain observed neural differences between BIPOC and non-BIPOC populations (68, 106).

The current study utilized a large population-based sample from the Adolescent Brain and Cognitive Development (ABCD) Study® to examine associations between perceived discrimination (PD) and the brain in childhood. Given similarities observed between childhood and adulthood, associations between experienced discrimination and amygdala connectivity to regions in the DMN (61), SN (52, 64), and thalamus (52; 55) were examined. We also evaluated the influence of behavioral and environmental factors on resulting associations. These relationships were also examined in a sub-sample of BIPOC children due to the increased rates of discrimination (21, 22) and experienced adversity (68, 106) observed in BIPOC populations. These findings may advance understanding of early mechanisms underlying negative health outcomes and inform the development of targeted interventions.

## 2. Methods and materials

### 2.1. Participants

Participants were drawn from the ongoing ABCD study, a large-scale study following 9- to 10-year-old children recruited from 21 sites across the United States (107). The study was approved by the Institutional Review Board at the University of California, San Diego. Informed consent was provided by parents and children. Data was accessed from the National Institutes of Mental Health Data Archives. A full sample of youth and (*N* = 4512) a subsample of BIPOC youth (*N* = 1567) was analyzed. Demographic data is provided in **Table S1** and **Table S2** in the Supplementary Materials. Only subjects without missing data, and with resting state data that met the criteria listed below, were included in our study.

### 2.2. Perceived Discrimination

Experienced discrimination was measured using the Perceived Discrimination Scale (108) at the one-year follow-up time point. Youth rated the extent to which they had experienced general discrimination over the past year across seven items on a five-point Likert scale, where higher scores indicated greater experienced discrimination.

### 2.3. MRI and fMRI data acquisition and processing

Resting-state functional magnetic resonance imaging (rsfMRI) data from the ABCD study from the baseline timepoint was sourced from NDA’s collection 3165-DCAN Labs ABCD-BIDS (https://nda-abcd-collection-3165.readthedocs.io/latest/pipelines/#abcd-bids-fmri/), using the ABCD downloader (https://github.com/DCAN-Labs/nda-abcd-s3-downloader). Image acquisition details are available in Hagler et al., 2019 (109). Data was processed through an adapted version of the HCP pipeline (110) provided by the DCAN Lab (https://github.com/DCAN-Labs/abcd-hcp-pipeline). Additional details can be found here: (https://nda-abcd-collection-3165.readthedocs.io/). After processing, to minimize head motion-related artifacts, a framewise displacement threshold of 0.2 mm was applied, and frames exceeding the threshold were removed (111). All data included in our analyses met the Data Analysis, Informatics & Resource Center’s quality control conditions (https://abcdstudy.org/study-sites/daic/). Analyses were restricted to participants with at least 8 minutes of usable resting-state data.

### 2.4. Resting-state amygdala functional connectivity

Left and right amygdala regions of interest (ROIs) were extracted from the Gordon Parcellation, which consists of 333 cortical parcels across 13 resting-state networks (see Fig. 1A, 60) in addition to 19 subcortical ROIs (see Fig. 1B) and cerebellum for a total of 352 ROIs grouped into 14 networks (112). Right and left amygdala ROIs were used as seed regions to correlate with all other regions from the Gordon parcellation, and correlation values between each amygdala ROI to all other ROIs were generated. Right and left amygdala connectivity was evaluated separately. In the current study, we chose to examine right and left amygdala connectivity to the DMN, SN, and thalamus based on previous literature indicating these connections may be impacted by experienced discrimination.

**Fig 1.**
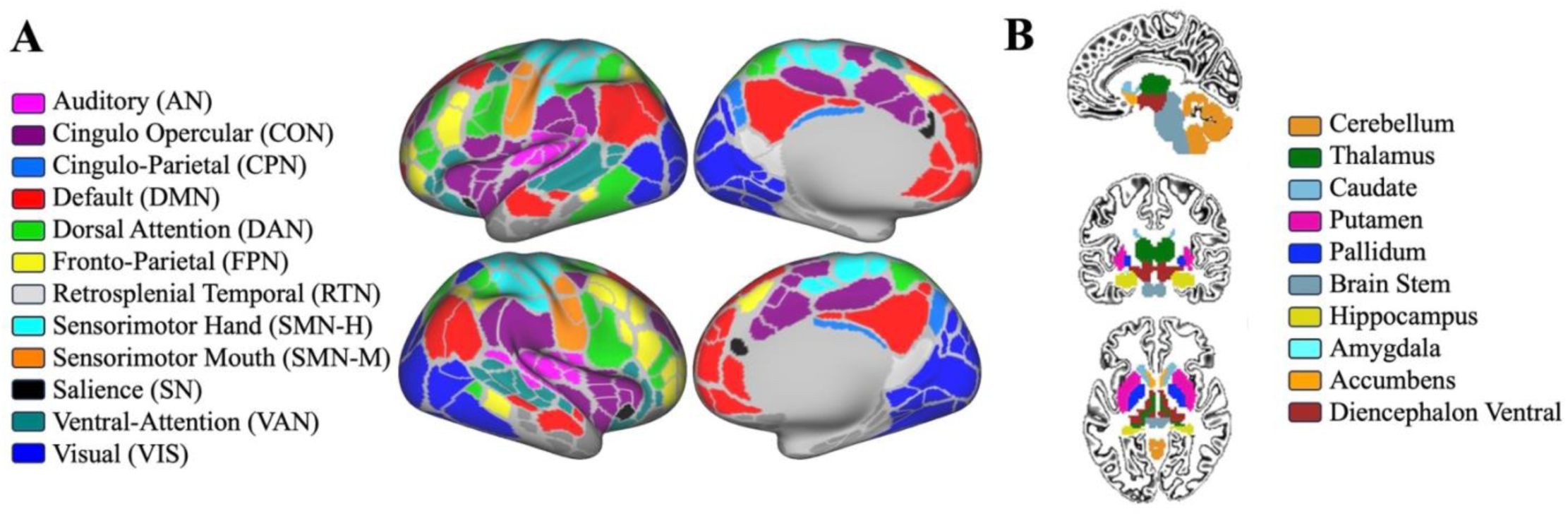
Cortical and subcortical parcellation schema **A.** Gordon Parcellation ROIs (60) color-coded by functional network. **B.** Subcortical volume ROIs.

## 3. Statistical analyses

Linear mixed-effects models were used to identify associations between amygdala connections and PD. We evaluated left and right amygdala connectivity in two separate models. Each model included connectivity to the right and left thalamus, and all nodes in the SN and DMN. Participants with any missing data were excluded from analyses. Sex, race/ethnicity, age, handedness, parental education, puberty status, and scanner ID were included as fixed effects, with family nested in scanner type included as a random effect. All resulting p-values were FDR corrected.

## 4. Results

### 4.1. Amygdala connectivity is associated with perceived discrimination

We examined the relationship between PD right and left amygdala connections to the SN, DMN, and right and left thalamus, as indicated above, from the full youth sample and BIPOC subsample. In our full sample, results showed a significant negative association between PD and right amygdala connectivity to the right thalamus (β = −0.02 and *p* < .05), indicating lower connectivity between the right amygdala and right thalamus was linked to greater experienced discrimination (see Fig. 2). Additionally, a negative trend-level association was observed between PD and right amygdala connectivity to the left thalamus (β = −0.01 and *p* < 0.1) such that lower connectivity was associated with greater experienced discrimination. Furthermore, a positive trend-level association was observed between PD and right amygdala connectivity to a DMN region (L ventromedial PFC [vmPFC], β = 0.02 and *p* < 0.1), such that greater connectivity was associated with greater experienced discrimination.

**Fig 2.**
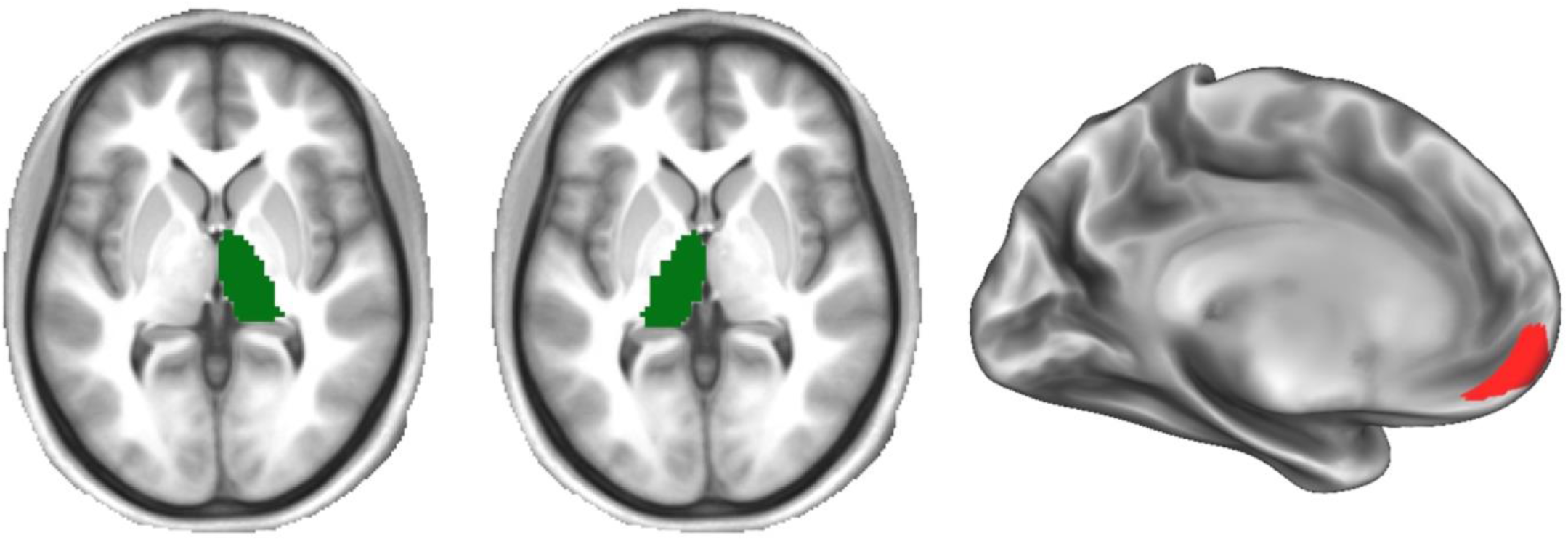
Right amygdala connections significantly associated with PD in the full sample. Colors represent subcortical thalamus (green) and cortical DMN (red) regions.

Within the BIPOC subsample, a positive trend-level association was found between PD and right amygdala connectivity with an SN node in the L ACC (β = 0.02, *p* < .1), indicating increased connectivity between the right amygdala and L ACC was linked to greater experienced discrimination (see Fig. 3). Additionally, a positive trend level association between PD and left amygdala connectivity to a DMN region (L angular gyrus, β = 0.02, *p* < .1) and a negative trend level association between left amygdala connectivity to the left thalamus (β = −.02, p < .1) were found. Thus, greater experienced discrimination was linked to increased left amygdala-L angular gyrus connectivity and decreased left amygdala-left thalamus connectivity (see Fig. 3).

**Fig 3.**
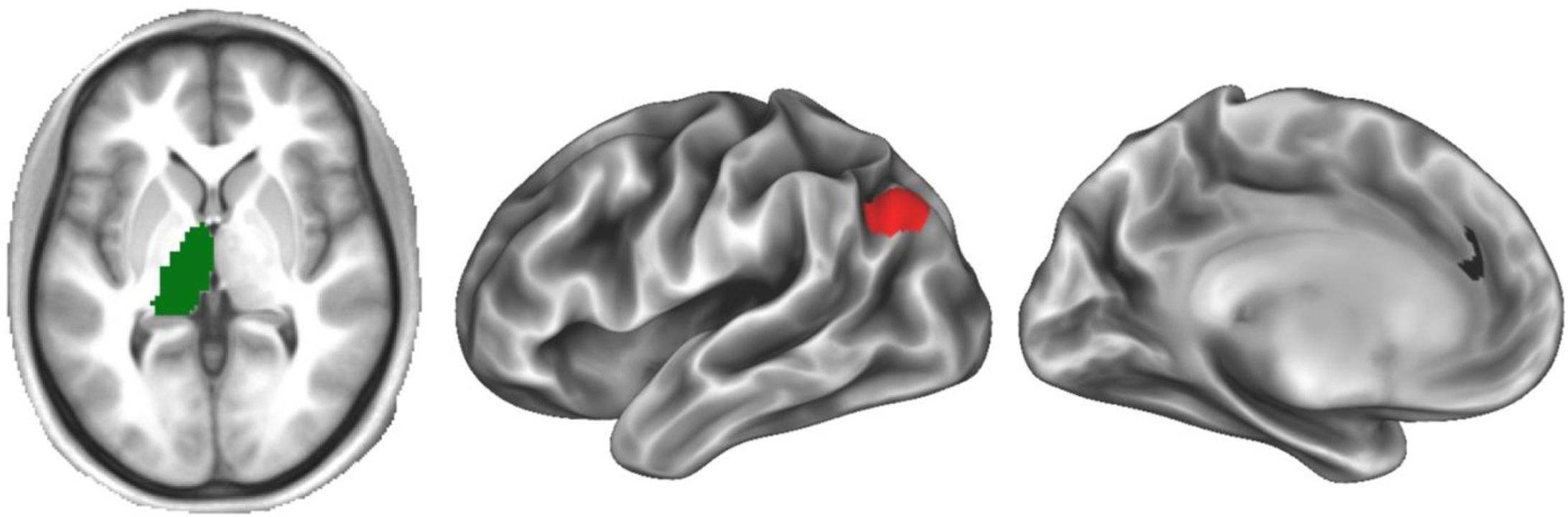
Amygdala connections associated with PD at a trend level (p < .1) in the BIPOC sample. Colors represent left amygdala-left thalamus (green) and left amygdala-left cortical DMN (red) and right amygdala-left SN (black) regions.

### 4.2. Traumatic events moderated the relationship between amygdala connections and perceived discrimination

Behavioral and environmental factors that may impact associations found between amygdala connectivity and PD were examined as potential moderators in a series of secondary analyses. Separate models examined internalizing, externalizing, traumatic events experienced, English proficiency, prosocial behavior, family environment, and school environment as potential moderators through the creation of interaction terms. Information about how these measures were collected can be found in our *Supplementary Materials*. Separate models were run for each amygdala connection with significant or trend-level (*p* < .1) associations with PD in both full and BIPOC samples. All analyses included sex, race/ethnicity, age, handedness, parental education, puberty status, and scanner ID as fixed effects with family nested in scanner type included as a random effect (FDR correction: *p* < .05). The following model was run for each variable to test the degree with which each variable moderated the relationship between each amygdala connection and PD:

Amygdala connectivity = intercept + *ԁ_1_* + β_1_(Sex) + β_2_(Race/Ethnicity) + β_3_(Age) + β_4_(Handedness) + β_5_(Parental Education) + β_6_(Puberty Status) + β_7_(Scanner ID) + β_8_(Main Effect) + β_9_(moderating variable x PD) + e,

where *ԁ_1_* represents random effects, e represents error, and β represents the weighted contribution of each variable.

Results indicated that traumatic events significantly moderated the association between amygdala connectivity to the left ACC and PD in our BIPOC subsample. Specifically, BIPOC children with higher experiences of traumatic events had stronger negative associations between experienced discrimination and right amygdala connectivity to the left ACC (β = −.03, *p* < .05). Other factors examined did not moderate associations between amygdala connectivity and PD.

## 5. Discussion

### 5.1. Summary of results

The current study examined associations between amygdala connectivity and PD in a full sample and BIPOC subsample of children. In a series of secondary analyses, the potential influence of behavioral and environmental factors on identified significant and trend-level associations was also examined. Our study revealed distinct relationships between amygdala connectivity and PD in the full sample versus BIPOC subsample of youth. In our full sample, stronger right amygdala connectivity to the right thalamus was significantly associated with lower levels of PD. Right amygdala connectivity to the left thalamus was also positively associated with PD at a trend level. However, in our BIPOC subsample, only trend-level associations were found. These showed positive associations between experienced discrimination and amygdala connectivity to a DMN region (angular gyrus) and SN region (ACC), and negative associations with amygdala connectivity to the thalamus. In this subsample, traumatic events significantly moderated the relationship between right amygdala-left ACC connectivity and PD, such that greater experienced trauma predicted stronger positive associations between this connection and PD. Our results suggest distinct amygdala connections relate to experienced discrimination in BIPOC children, where experiences of traumatic events may further impact observed effects on this population. Broadly, our results suggest discrimination may have unique effects on minoritized populations and may impact neural connections associated with risk for psychopathology.

### 5.2. Amygdala connectivity to thalamus linked to discrimination in full sample

In our full sample, greater PD was significantly linked to decreased right amygdala connectivity to the right thalamus (*p* < .05) and left thalamus at a trend level (*p* < .1). This association contrasts with prior literature in adults, which found positive associations between this connection and experienced discrimination (52; 55). However, this connectivity pattern has been associated with threat responsivity (113, 114) and increased PTSD symptomatology (115) in adults. Prior studies also suggest experiences of discrimination may heighten PTSD symptoms (116–118), which may indicate that children with this connectivity pattern are at risk for developing these symptoms.

In our full sample, greater PD was also linked to increased amygdala connectivity to the vmPFC, a node in the DMN implicated in emotion regulation in adulthood (119) and childhood (120). In adulthood, connectivity changes in this network have been linked to experienced discrimination (56, 61), though this particular node has not been implicated in this context. Increased synchrony between the medial prefrontal cortex, more broadly, and basolateral amygdala has been associated with the ability to identify threat vs. safety cues (121). This suggests children with this connectivity pattern may be more likely to perceive instances of discrimination.

### 5.3. Amygdala connectivity not significantly associated with discrimination in BIPOC sample

In our BIPOC sub-sample, trend-level associations were found between amygdala connectivity and PD. Greater experienced discrimination predicted stronger amygdala connectivity to a DMN region (angular gyrus) and SN region (ACC), and lower amygdala connectivity to the thalamus. In line with prior research, the SN and DMN have been significantly associated with discriminatory experiences (55, 61) in BIPOC adults. Similar to results in our full sample, these patterns may relate to alterations in threat response and affective processing (49, 113, 114, 122). Unlike our full sample, we did not see a significant association between amygdala-thalamus connectivity and experienced discrimination, though a similar trend-level association was found, where greater connectivity was linked to lower discrimination. Our lack of significant findings may be explained by reduced statistical power, as our BIPOC sample size did not meet the recommended minimum for brain-wide association studies (123). Decreased between-group variance and the restricted range of discrimination scores within the BIPOC group may also explain the absence of significant associations.

### 5.4 Traumatic events moderated association between amygdala-ACC connectivity and discrimination in BIPOC sample

In a series of secondary analyses, behavioral (internalizing and externalizing) and environmental (school environment, prosocial behavior, English proficiency, traumatic events, and family conflict) factors were considered as potential moderators.

In our BIPOC sub-sample, experienced traumatic events moderated the relationship between amygdala connectivity to the ACC and PD, indicating that positive associations between discrimination and amygdala-ACC coupling strengthened as traumatic events experienced increased. The ACC and amygdala have often been implicated in PTSD (124–126). Increased connectivity between these structures has been associated with greater PTSD symptoms in Black adults (125), suggesting experiences of discrimination increase risk for developing these symptoms (116) in BIPOC individuals. Our results may indicate that discrimination-related stress in BIPOC children contributes to cumulative trauma, resulting in greater impacts on the brain, especially in regions relevant for emotional regulation and salience detection.

While not significant moderators, significant correlations were observed between PD and our environmental and behavioral factors (see **Table S2** and **Table S3)**. Significant positive correlations were seen between PD and both internalizing and externalizing behaviors in both samples, such that individuals with more experienced discrimination exhibited greater internalizing and externalizing behaviors. While associations between amygdala connectivity and internalizing (127–129) and externalizing (77, 130, 131) have been established in many studies, connectivity may be linked to specific subtypes of these behaviors as opposed to the overall constructs (79).

From the analyzed environmental variables, significant negative correlations were found between PD and school environment, prosocial behavior, and English proficiency in both samples. In line with prior literature (82), this may suggest that individuals in more supportive school environments exhibit more prosocial behavior, which facilitates social connection and resilience. Further, individuals with greater English language proficiency may experience less discrimination (98–100). Additionally, family conflict was positively correlated with PD, suggesting increased conflict in the home was related to greater experienced discrimination. Interrelations between school environment, prosocial behavior, language proficiency, and family conflict, and lack of variability within these measures, may explain why these factors did not individually moderate relationships between PD and the brain.

## 6. Limitations/Future directions

While the current study provides insight into the associations between amygdala connectivity and PD, several limitations must be acknowledged. First, relatively low variability in discrimination scores and our tested moderators may have hindered our ability to detect significant associations. Further, while experienced discrimination can be measured in childhood (27–29), our measure was based on the ABCD Study’s race and ethnicity classification (132) recommended by the NIH at the time of data collection, which may not have been the best fit for this time point (133). Additionally, because our BIPOC subsample was created by excluding participants categorized as white, individuals of Arab, Middle Eastern, and North African descent, who may identify as people of color, were likely not included. Multiracial individuals were also not considered, as no multiracial option was provided in our racial and ethnic classification variable. Future studies should expand racial and ethnic classifications and prioritize recruitment of diverse populations to better examine associations between discrimination and health outcomes (134). The relationship between discrimination and brain connectivity may have been impacted by other factors not evaluated in the current study, such as additional chronic stressors, coping strategies, and social support. Future studies should consider such factors. Lastly, this study lacks in its ability to assess longitudinal alterations in neural connectivity related to PD over time. Future research should address associations between amygdala connectivity and PD using future longitudinal data from the ABCD Study.

Our study underscores the impact of experienced discrimination on neural connectivity and emphasizes the importance of considering these experiences when examining health in youth, particularly in BIPOC individuals who may face compounded stressors. Understanding associations between the brain and these experiences provides a foundation for developing targeted interventions to mitigate the effects of discrimination on future health outcomes.

## Acknowledgments

Data used in the preparation of this article were obtained from the Adolescent Brain Cognitive Development (ABCD) study (https://abcdstudy.org), a multisite, longitudinal study designed to recruit more than 10,000 children aged 9-10 and follow them over 10 years into early childhood. The ABCD study is supported by the National Institutes of Health and additional federal partners under award numbers U01DA041048, U01DA050989, U01DA051016, U01DA041022, U01DA051018, U01DA051037, U01DA050987, U01DA041174, U01DA041106, U01DA041117, U01DA041028, U01DA041134, U01DA050988, U01DA051039, U01DA041156, U01DA041025, U01DA041120, U01DA051038, U01DA041148, U01DA041093, U01DA041089, U24DA041123, U24DA041147. A full list of supporters is available at https://abcdstudy.org/federal-partners.html. Information about participating sites and study investigators can be found at https://abcdstudy.org/consortium_members/. ABCD consortium investigators designed and implemented the study and/or provided data but did not necessarily participate in the analysis or writing of this report. The manuscript reflects the views of the authors and does not necessarily represent the opinions or views of NIH or ABCD consortium investigators. The ABCD data repository is dynamic and continues to grow over time. The research was supported by funding from Earlham College, including Alumni Council Fund for Faculty Research, the Department of Education’s Ronald E. McNair Postbaccalaureate Achievement Program, and the CBB-Student Faculty Research Support Fund.

