## Supplement for "Associations between amygdala connectivity and experienced discrimination in children"

**Supplementary material**

*Table S1.* Demographics for study participants.

| | | Full Sample N = 4512 | BIPOC Sample N = 1567 |
|---|---|---|---|
| **Sex (%)** | Male | 54.17 | 52.65 |
| | Female | 45.83 | 47.35 |
| **Handedness (%)** | Right | 80.81 | 81.43 |
| | Left | 6.98 | 6.51 |
| | Mixed | 12.21 | 12.06 |
| **Highest Parent Education (%)** | Less than HS | 1.84 | 4.60 |
| | HS or GED | 4.83 | 10.47 |
| | Some college | 9.97 | 16.15 |
| | Associate’s | 11.41 | 15.76 |
| | Bachelor's | 29.23 | 22.85 |
| | Master's | 28.48 | 20.23 |
| | Professional | 6.69 | 4.53 |
| | Doctoral | 7.54 | 5.42 |
| **Race/Ethnicity (%)** | White | 64.14 | — |
| | Black | 7.76 | 21.83 |
| | Hispanic | 16.49 | 45.25 |
| | Asian | 1.91 | 5.36 |
| | Other | 9.71 | 27.57 |
| **Puberty Scale (%)** | Prepuberty | 19.23 | 24.06 |
| | Early Puberty | 40.03 | 37.01 |
| | Mid Puberty | 29.12 | 36.50 |
| | Late Puberty | 1.51 | 2.17 |
| | Post Puberty | 0.11 | 0.26 |

| | | |
|---|---|---|
| **Age (months), Mean (SD)** | 120.09 (7.48) | 119.80 (7.54) |

*Table S2.* Pearson's correlations for full sample moderator variables ($N = 4408$)

| | **1** | **2** | **3** | **4** | **5** | **6** | **7** | **8** |
|---|---|---|---|---|---|---|---|---|
| **1. Perceived Discrimination** | — | | | | | | | |
| **2. Internalizing** | .09*** | — | | | | | | |
| **3. Externalizing** | .12*** | .55*** | — | | | | | |
| **4. Traumatic Events** | .08*** | .12*** | .15*** | — | | | | |
| **5. English Proficiency** | -.09*** | -.03 | .00 | .02 | — | | | |
| **6. Prosocial Behavior** | -.11*** | -.08*** | -.15*** | -.01 | .10*** | — | | |
| **7. Family Conflict** | .21*** | .08*** | .20*** | .04* | -.10*** | -.24*** | — | |
| **8. School Environment** | -.18*** | -.13*** | -.14*** | -.04** | .10*** | .35*** | -.23*** | — |

*$p < .05$, **$p<.01$, ***$p < .001$

*Table S3.* Pearson's correlations for BIPOC sample moderator variables ($N = 1567$)

| | **1** | **2** | **3** | **4** | **5** | **6** | **7** | **8** |
|---|---|---|---|---|---|---|---|---|
| **1. Perceived Discrimination** | — | | | | | | | |
| **2. Internalizing** | .09*** | — | | | | | | |
| **3. Externalizing** | .13*** | .58*** | — | | | | | |
| **4. Traumatic Events** | .05* | .13*** | .18*** | — | | | | |
| **5. English Proficiency** | -.10*** | -.03 | .03 | .04 | — | | | |
| **6. Prosocial Behavior** | -.08** | -.05 | -.10*** | .01 | .06** | — | | |
| **7. Family Conflict** | .22*** | .06* | .14*** | .04 | -.09*** | -.20*** | — | |
| **8. School Environment** | -.17*** | -.13*** | -.14*** | -.04 | .05* | .35*** | -.24*** | — |

*$p < .05$, **$p<.01$, ***$p < .001$

### Additional Behavioral and Environmental Factors

#### Internalizing and externalizing behaviors

Parent-reported measures of internalizing and externalizing behavior were obtained using the Child Behavior Checklist (CBCL; age 6 to 18 form) (Achenbach & Rescorla, 2001). Specifically, parents rated behaviors based on how characteristic they were of their child over the past six months. Items were combined to create composite scores for both internalizing and externalizing behavior, with higher scores associated with increased internalizing or externalizing behaviors.

#### Traumatic events experienced

Parent-reported traumatic events were assessed as part of the post-traumatic stress disorder module of the Kiddie Schedule for Affective Disorders and Schizophrenia (KSADS-5) (Kaufman et al., 1997). This scale contains 17 items asking about specific traumatic events, and participants rated whether or not stated events had been experienced by their child (0 - no, 1 - yes) to assess the number of traumatic events experienced.

#### English proficiency

Child-reported measures of English-language proficiency were obtained using the PhenX Youth Acculturation Survey (Hamilton et al., 2011). Specifically, youth rated their fluency in English across seven items related to which language(s) they use daily. Specifically, we analyzed question one, which asked participants how well they speak English, where higher scores were associated with increased English-speaking proficiency.

#### Prosocial behavior

Child-reported measures of prosocial behavior were obtained using the Prosocial Behavior Survey, a shortened three-item variable from the Strength and Difficulties Questionnaire Prosocial Scale (Goodman et al., 1998). Youth reported how they interacted with others to measure prosocial interactions, with increased scores associated with increased prosocial behavior.

#### Family environment - conflict

Child-reported measures of conflict within their family environment were measured with the PhenX Family Environment Scale (Hamilton et al., 2011). Youth rated whether or not nine items that assessed the quality of interpersonal relationships within the family were characteristic of their family.

#### School environment

Child-reported measures of school environment were measured with the PhenX School Risk & Protective Factors Survey (Hamilton et al., 2011). Specifically, youth rated how much they agreed with six statements about a supportive school environment, where increased scores were associated with more supportive school environments.

*Figure S1.* Traumatic events experienced as a moderator

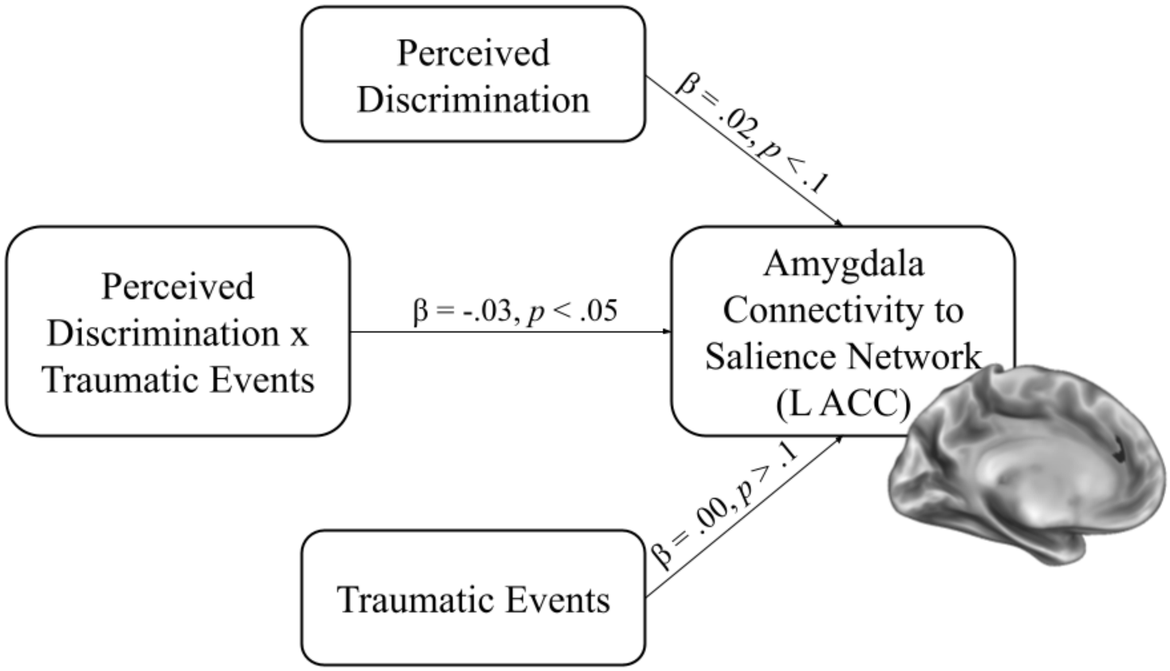


*Figure S2.* Simple slopes analysis considering traumatic events experienced as a moderator

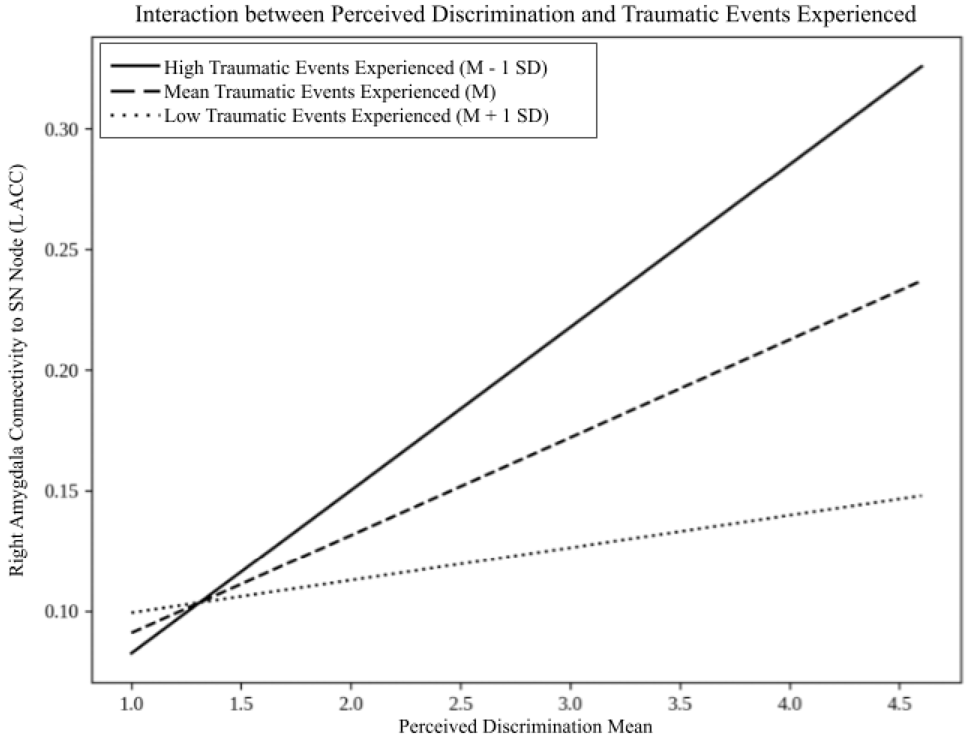
