## Supplementary figures and images for "Associations between amygdala connectivity and experienced discrimination in children"

### Supplement Figure 1

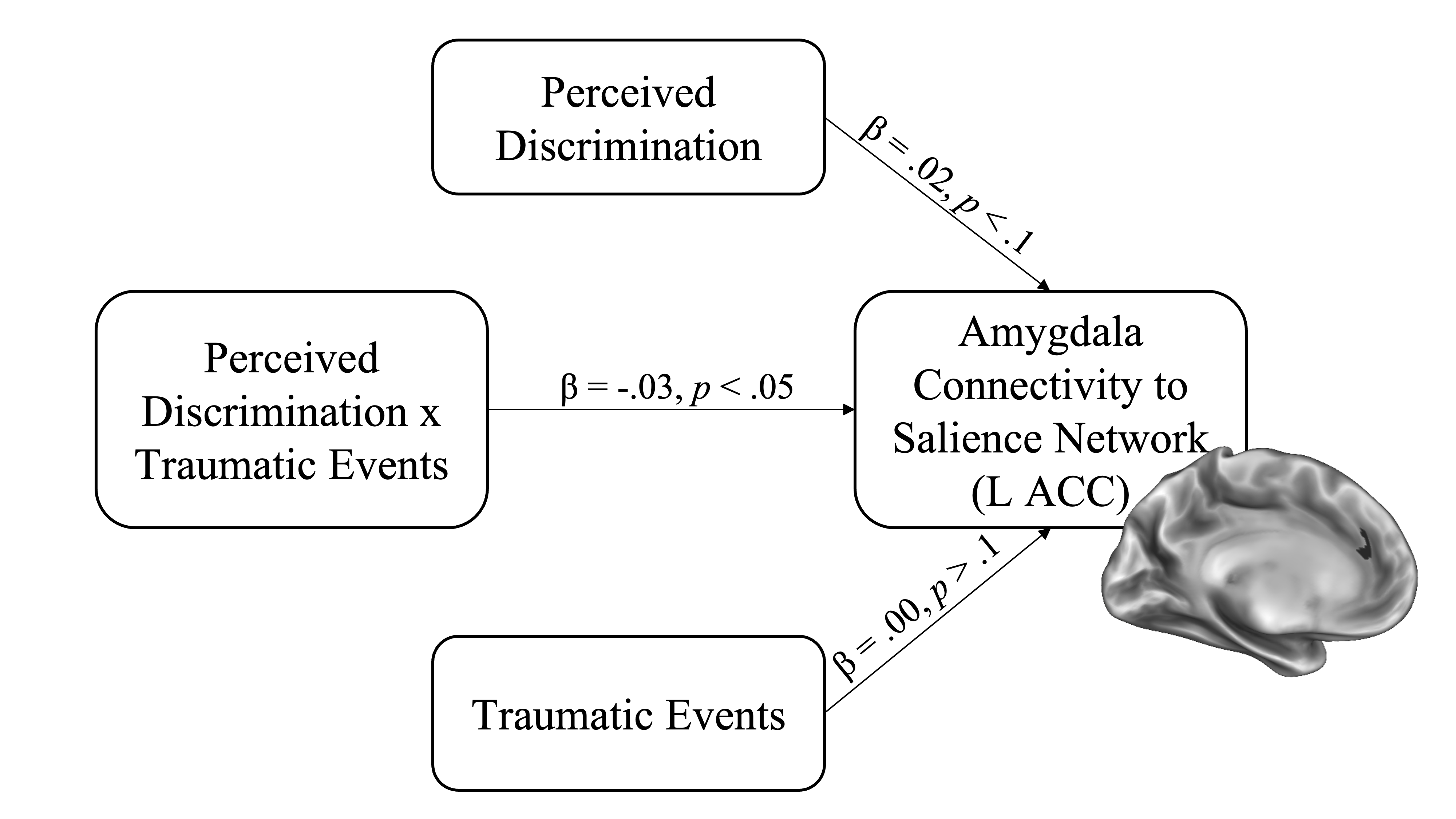

### Supplement Figure 2

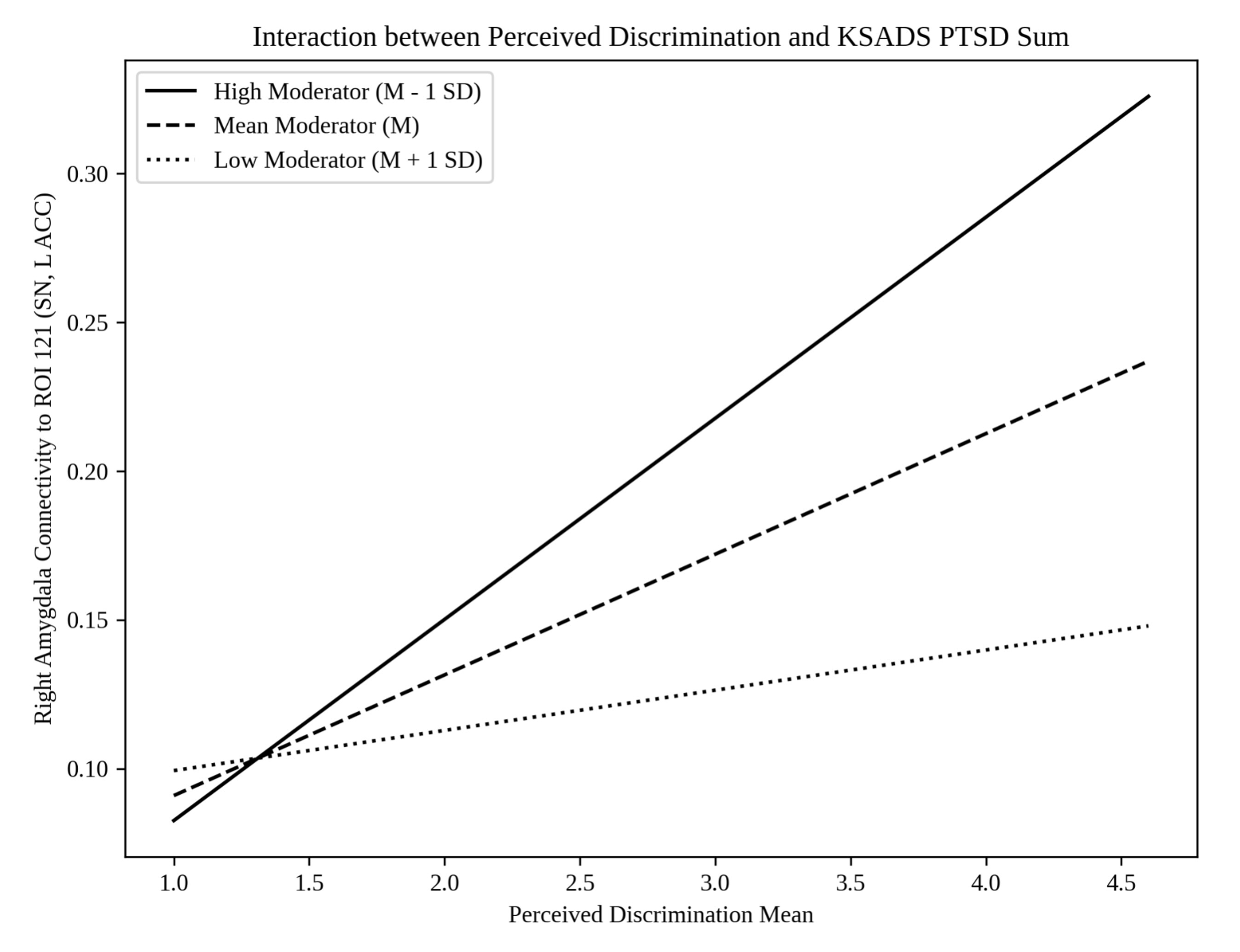
